# Peptide-DNA Conjugates for Formation of Lipid Nanodiscs

**DOI:** 10.64898/2026.05.14.725262

**Authors:** Praneetha Sundar Prakash, Soumya Chandrasekhar, Joy Kabuga, Diana P. N. Gonçalves, Fatemeh Fadaei, Thorsten-Lars Schmidt

## Abstract

Nanoscale lipid bilayer mimetics are powerful tools for research on lipid bilayer, membrane proteins or for drug delivery. Established nanoscale bilayer systems that are stabilized by short peptides or polymers produce a broad size distribution and are difficult to customize. Here we introduce a DNA nanotechnology-based lipid bilayer mimetic, in which we covalently conjugated established nanodisc-forming amphiphilic peptides to oligonucleotides. These peptide-DNA conjugates were then hybridized with a circular single-stranded scaffold to form stiff, circular PDC minicircles with 14 peptide modifications at the inner rim of the torus. Lipid reconstitution yielded defined nanodisc with a tightly controlled circumference and component stoichiometry. Molecular dynamics simulations further validated the structural stability and reveal an asymmetric migration of the DNA to one rim of the bilayer. To mimic membrane protein insertion, we co-reconstituted a transmembrane peptide coupled to a bulky quantum dot. In future applications, the size and peptide arrangement can easily be modified in these DNA-templated PDC nanodiscs.

---

About 30% of the human genome codes for membrane proteins (MPs) and contain highly hydrophobic domains that anchor to cellular lipid membranes. MPs play important roles in many cellular functions such as environment sensing, intercellular communication and, substrate attachment and are the target for most commercially available drugs.^1^ Yet, only a small fraction of unique solved structures in the Protein Data Bank are membrane proteins,^2^ which can be attributed to the experimentally complex expression and purification in practical quantities.^3^ Moreover, hydrophobic domains promote aggregation of MPS which inhibits their structural characterization by X-ray crystallography or Cryo-Electron microscopy (EM).^4^ Detergents can solubilize MPs but often alter their structural conformation of proteins^4,5^ affect the buffer viscosity, surface tension, vitrified ice thickness, particle distribution within the ice and are therefore problematic in structure determination by cryo-EM.^6–8^

Apolipoprotein-derived membrane scaffolding protein (MSP) nanodiscs^9,10^ allow solving membrane protein structures in their lipid environments by co-reconstitution of detergent-solubilized membrane proteins, lipids and MSPs. The sizes of MSP nanodiscs can be controlled by the length of the MSP protein, and covalent circularization of the MSP nanodiscs can narrow size distribution and form larger nanodisc complexes. However, any design modification requires elaborate protein engineering and expression.^10–12^

Other popular bilayer mimetics include amphiphilic saposin-lipoprotein (Salipro) nanodiscs,^13^ styrene maleic acid (SMA) copolymers-based nanodiscs^14^ or peptide-based nanodiscs or “peptidiscs”.^15,16^ The peptidiscs-forming peptides are derived from one alpha-helical segment of the backbone of MSP proteins and can be synthesized chemically by solid phase synthesis.

Peptidiscs are useful bilayer mimetics for biophysical studies of membrane proteins with Nuclear Magnetic Resonance (NMR) or cryo-EM.^15,16^ The size distribution of all these nanodiscs is not as narrow as for MSP nanodiscs and depends on the stoichiometric ratio between the amphiphilic molecule to lipid.

To achieve better size control and to be able to introduce modifications in a programmable way, we developed DNA-encircled lipid nanodiscs.^17^ In this DNA nanotechnology-based system, a lipid bilayer is reconstituted inside a modified DNA minicircle. The interface between lipids and the DNA circle was formed by hydrophobic S-alkyl modifications of phosphorothioate groups in the backbone.^17^ While coarse-grained molecular dynamics (MD) simulations suggested that increasing the number of alkyl groups would form a tighter interface and form more stable nanodiscs,^18^ we recently found that excessive S-alkylation reduces thermal and structural stability of the DNA.^19^ We are therefore developing alternative DNA modifications to stabilize the lipid bilayer inside the DNA ring. In an alternative approach, we modified DNA with amphiphilic polyethylene glycol (PEG) molecules, to interface with the lipid bilayer.^20^ While the PEG modification did not significantly reduce the stability of the DNA, this approach still required cationic lipids and detergents during the synthesis, which can destabilize membranes and denature membrane proteins.^21^

Herein, we explore the modification of the DNA oligonucleotides with peptidiscs-forming peptides for the synthesis of nanodiscs (Figure 1). For this, we first synthesized peptide-DNA conjugates (PDC) from short oligonucleotides and peptides, and hybridized multiple copies of PDC oligonucleotides to a circular single-stranded (ss) DNA circles to form PDC minicircles. Other than in peptidiscs, the size of peptide-modified DNA circles and the number of peptides per complex can be precisely controlled through the DNA design. For the reconstitution of lipids, cationic detergents or lipids are not required anymore as in previous DNA-based designs,^17,20^ expanding their potential in membrane protein research.

**Figure 1.**
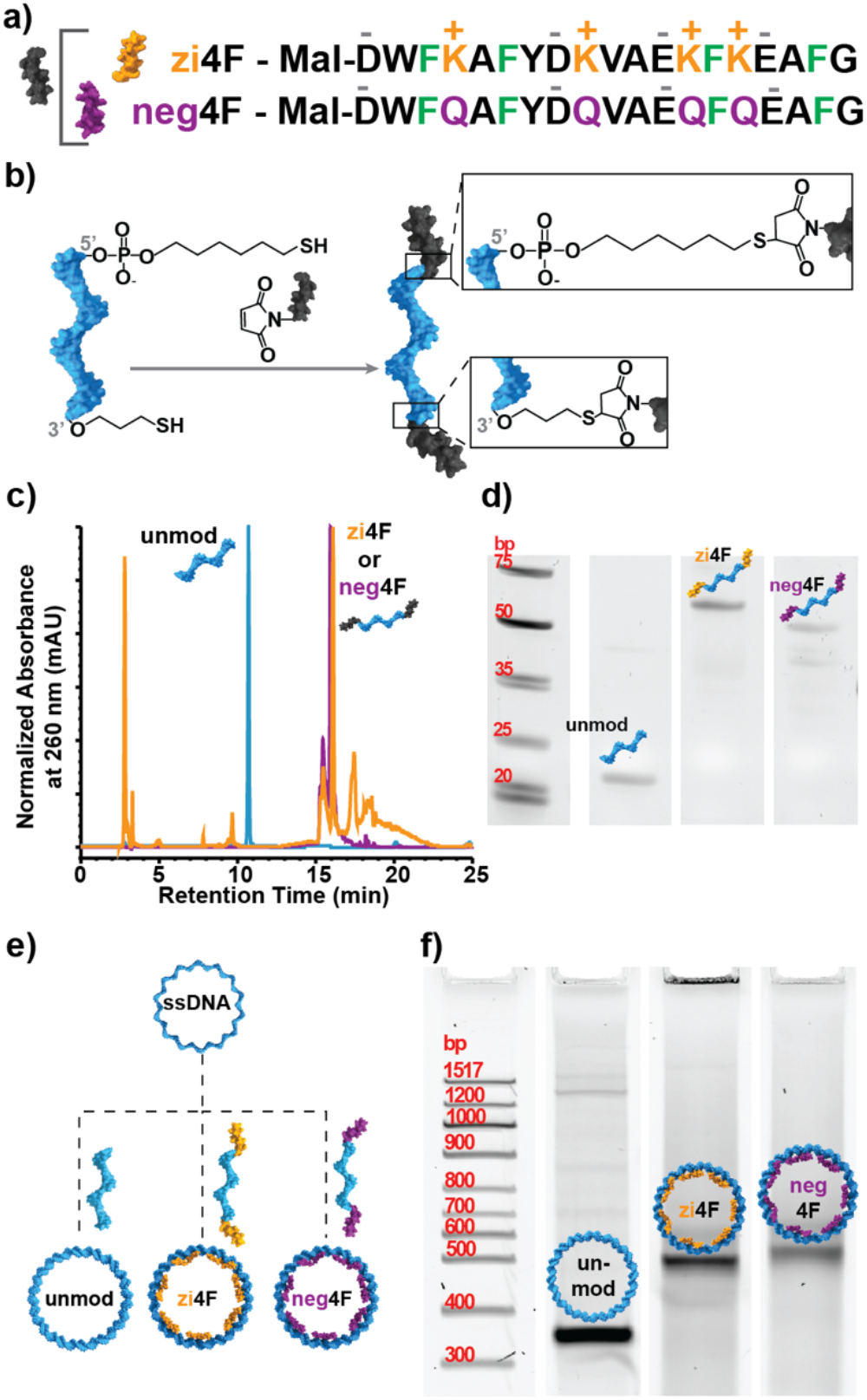
Formation of peptide-DNA conjugates (PDCs) and minicircles. a) The two different peptides used in this study: the zwitterionic zi4F and the modified, negatively charged neg4F peptide; Both carry a thiol-reactive maleimide (mal) at the N-terminus. b) Formation of the peptide-DNA conjugate (PDC) oligonucleotides. c) Reverse phase HPLC chromatogram of the unmodified oligonucleotide and the PDC oligonucleotides with two neg4F or zi4F peptides. d) 10% Denaturing urea PAGE gel of the oligonucleotide with two thiol-modifications and the two different PDCs. e) The oligonucleotides were then hybridized to covalently circularized single stranded DNA to form double-stranded (ds) minicircles. f) 5% native PAGE of unmodified or PDC modified dsminicircles.

## Results and Discussions

### Synthesis of PDC Conjugates

To stabilize the lipid bilayer in the DNA rings, we explored two different amphiphilic peptide sequences (Figure 1a): the zwitterionic 4F peptide (zi4F) and the negatively charged “neg4F” peptide. Both peptides are alpha helices with ∼3.6 residues per turn with a hydrophilic and a hydrophobic side. The zi4F peptide contains four anionic amino acids (D = aspartic acid, E = glutamic acid) and four cationic lysines (K), while four phenylalanines (4F) and one tryptophan (W) are found on the hydrophobic side. In the neg4F sequence, we exchanged all cationic lysines for neutral, hydrophilic glutamines (Q). Both peptides were synthesized with maleimides at the N-terminus and were conjugated to the 3’ and 5’ ends of thiol-modified oligonucleotides Figure 1b).

The neg4F peptide was designed because the zi4F peptides tended to aggregate with DNA at a high molar excess during conjugation or when assembled into double-stranded minicircles (Figure 1f), which we attributed to electrostatic interactions between their lysines and phosphates of the oligonucleotides. The modified neg4F peptides do not contain any cationic residues and do not aggregate with DNA.

In reverse phase high pressure liquid chromatography (RP-HPLC), the oligonucleotide-peptide conjugates (Figure 1c) eluted slower than unmodified control oligonucleotides due to the added hydrophobicity, in the peptides. The purified conjugates also migrated slower than unmodified oligonucleotides in denaturing polyacrylamide gel electrophoresis (PAGE, Figure 1d) proving successful conjugation.

### Formation of PDC Minicircles

To assemble double-stranded (ds) minicircles, a single-stranded (ss) DNA minicircle template was synthesized with seven repeats of a sequence that is complementary to the short oligonucleotides by enzymatic splint ligation (Table S 1 and Figure S 1). The ss minicircle template was then hybridized with the short oligonucleotides (unmodified or zi4F/neg4F PDC) to form unmodified or PDC minicircles respectively (Figure 1e and Figure 2). In native PAGE, ds PDC minicircles also migrate slower than the unmodified ds minicircles (Figure 1f). They were also characterized using TEM and AFM imaging and exhibited narrow size distribution between 17±2 nm (Figure 2 and Figure S 2).

**Figure 2.**
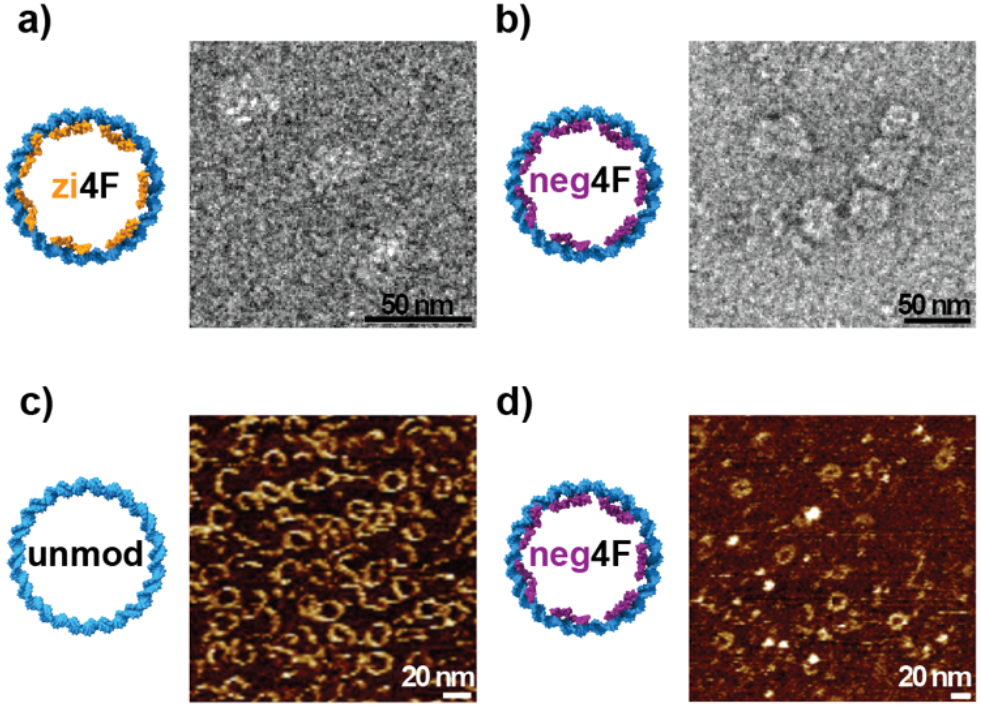
TEM and AFM micrographs of DNA minicircles. a-b) TEM micrographs of zi4F and neg4F minicircles. Due to staining artefacts, the empty minicircles appear to be filled. c-d) AFM images of unmodified and neg4F minicircles clearly indicate that the interior of these mini-circles is hollow.

In templated DNA nanotechnology including DNA origami,^22^ a high molar excess (often 10fold) of short oligonucleotides (staples) is used over the templates to maximize incorporation probability of short staples. Previously, we used molecular weight cut-off filters for the purification of the ds minicircles from excess of short oligonucleotides as these might impede the formation of nanodiscs. For ds-PDC minicircles, the purification yields were however poor (<10 %), presumably due to non-specific binding of the conjugates to the filter membranes. Filter passivation with Pluronic™ increases recovery yields of DNA structures after ultrafiltration,^23^ but was not tested here as Pluronic™ is a surfactant and traces in the sample could destabilize lipid nanodiscs in the following reconstitution step. PDC minicircles however also formed well with a modest 1.5-fold excess of PDC oligonucleotides per binding site (or 10.5-fold molar excess over scaffold) and ultrafiltration was omitted (Figure S 3).

While neg4F PDC minicircles were stable, the zi4F PDC minicircles aggregated at higher concentrations (Figure S 4), which we attributed to electrostatic interactions of the cationic lysines with the negative phosphate backbone of DNA. This was further confirmed through STEM images (Figure S 2e) where the zi4F PDC minicircles appeared to be collapsed (Figure S 2c) while unmodified and neg4F PDC minicircles remained circular (Figure S 2f).

Finally, we tested which detergents are compatible with neg4F minicircles during lipid reconstitution and identified sodium cholate and CHAPS as the best candidates, since they did not aggregate the minicircles (Figure 3 and Figure S 5).

### Reconstitution of Lipids

To reconstitute a lipid bilayer into the hydrophobic center of PDC minicircles, they were incubated with detergent-solubilized lipids and detergents were removed by passing the sample through detergent removal spin columns. PDC nanodiscs formed with both detergents (sodium cholate and CHAPS) and with both zi4F and neg4F PDC rings. The formation of PDC nanodiscs was confirmed by gel electrophoresis and TEM imaging (Figure 3).

**Figure 3.**
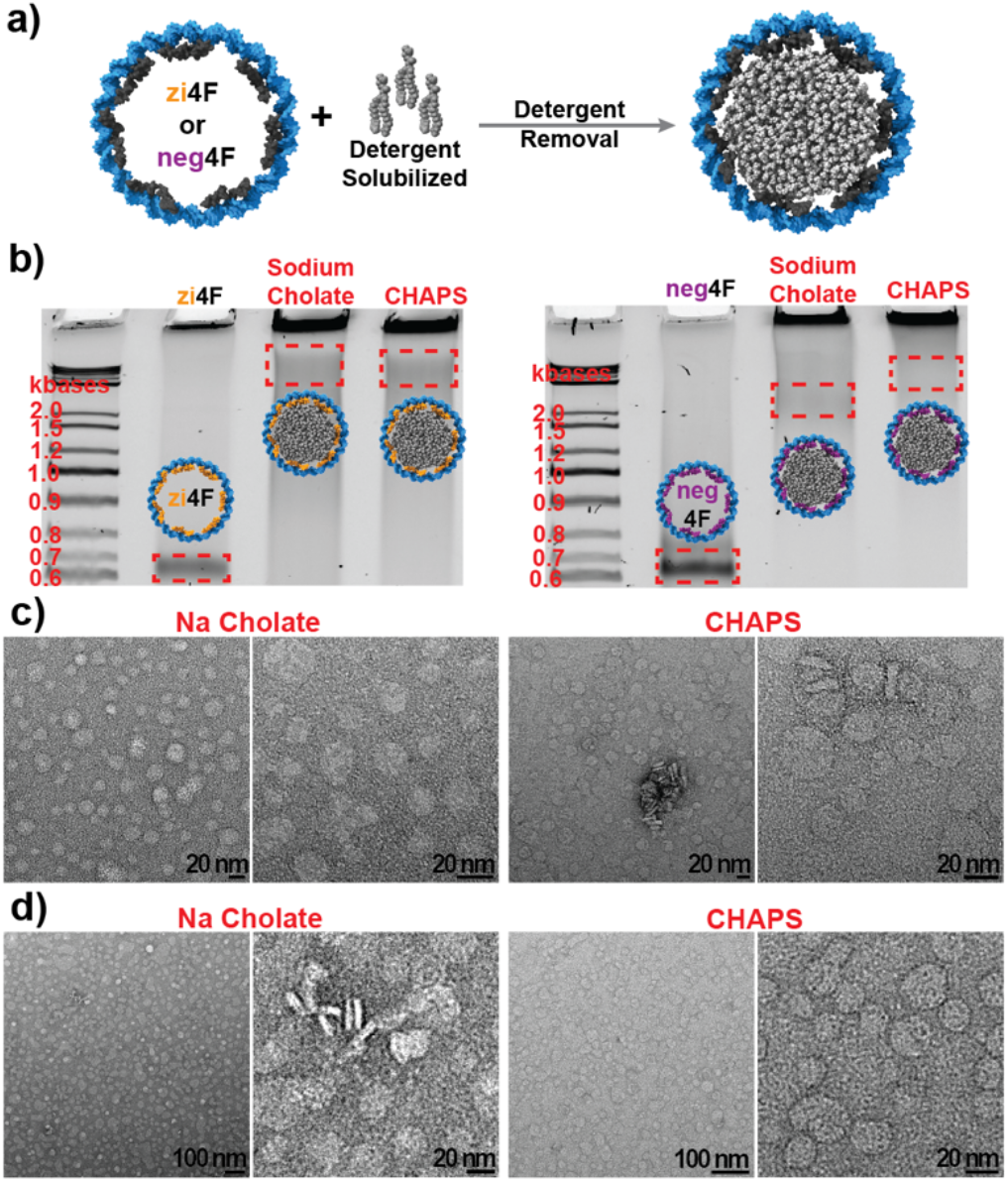
Formation of PDC lipid nanodiscs. a-b) Schematic representation of the reconstitution process and corresponding native PAGE gel analysis of the complexes in a) zi-4F PDC minicircles and b) neg4F PDC minicircles and the corresponding native PAGE gels. DMPC lipids were added as detergent-solubilized mixed micelles in sodium cholate or CHAPS as indicated. c-d) corresponding TEM images.

Our previous DNA-lipid nanodiscs systems^17,20^ could not be analyzed by native PAGE due to cationic lipids and detergents. PDC lipid nanodiscs on the other hand formed a broad band with a lower electrophoretic mobility than empty PDC minicircles (Figure 3a-b). While the gel lane shows a strong hanging lane, which possibly corresponds to nanodisc-aggregates and/or liposomes which did not interact with the peptides, there also appears a sizeable population of formed nanodiscs. Note that the empty ds neg4F does not show any aggregation.

Samples were also drop-cast on TEM grids and imaged directly after detergent removal, revealing nanodiscs with a narrow size distribution around 18±3 nm (Figure 3c-d). PDC lipid nanodiscs (Figure 3c and d) had better contrast and clearer defined boundaries than empty PDC minicircles (Figure 2 and Figure S 2). A small population of larger nanodiscs (>20 nm) could be caused by scaffold dimers that are a side product from the enzymatic splint ligation^17^ or fusing of neighboring nanodiscs during drying of the grids.

In size exclusion chromatography (SEC), zi4F PDC conjugates were again more problematic than the neg4F versions and did not elute due to non-specific sticking to either the column filter or resin. There was no significant elution volume shift between neg4F PDC minicircles and nanodiscs (Figure S 6). This could be explained by the similarity of the hydrodynamic radius of both complexes. Other than nanodisc-forming MSP proteins that are flexible and collapse without lipids, circularized DNA is very stiff and circles do not supercoil in this size range.^24,25^

The PDC nanodisc peak was collected after SEC, concentrated and imaged by TEM, showing a mix of nanodiscs and liposomes. We hypothesized that the formation of liposomes might be caused by an insufficient shielding of the hydrophobic rim of the lipid bilayer by the peptides and DNA.

### Molecular Dynamics Simulations

To better understand the nature and dynamics of the lipid– peptide–DNA interface, we performed 200ns all-atom molecular dynamics simulations of both zi4F (Movie S 1 and S 2) and neg4F (Movie S 3 and S 4) peptide–DNA conjugate (PDC) nano-discs. For this, models of both systems were constructed using the same DNA sequence and oligonucleotide functionalizations as in the experiments. Figure 4 shows a side view of the functionalized DNA nanodiscs over the 200 ns simulation. While both peptide chains in each system were initially aligned with the DNA, their orientation shifted over time. Close-up views (Figure 4c and Figure 4d) illustrate stable interactions between the residues of the peptide and the lipid core. Specifically, these aromatic rings orient toward the lipids to establish and maintain the stability of the systems throughout the simulation. To characterize peptide–lipid interactions and structural behavior, we analyzed residue-specific distances to the lipid bilayer (Figure S 7) as well as the time evolution of peptide helix orientation (Figure S 8).

**Figure 4.**
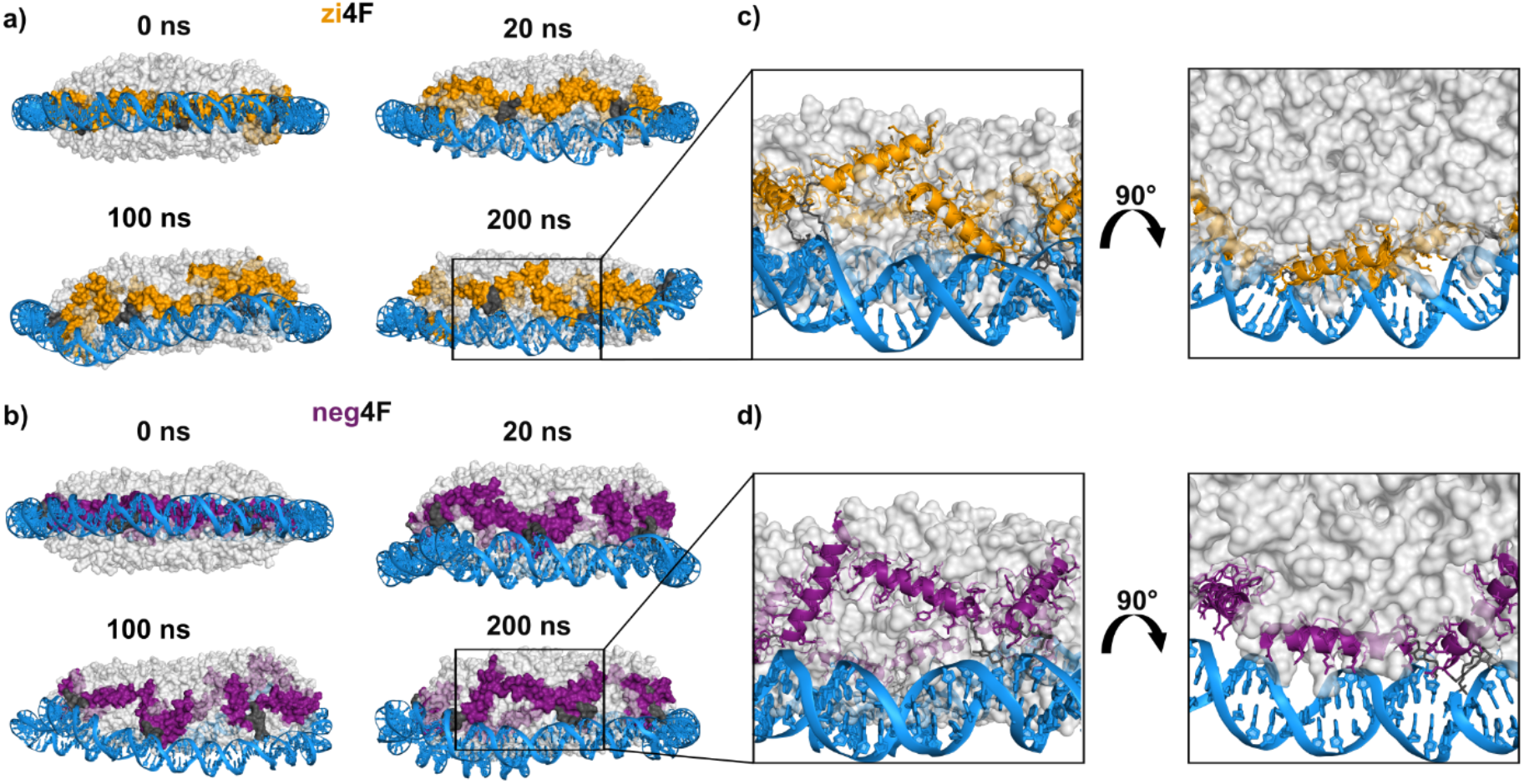
MD snapshots of DNA nanodiscs functionalized with zi4F and neg4F peptides. (a,b) Time evolution (0–200 ns) for zi4F (a) and neg4F (b). (c,d) Enlarged 200 ns views with corresponding 90° rotated orientations for zi4F (c) and neg4F (d). DNA (blue), peptides (orange for zi4F, purple for neg4F), lipids (gray), and linkers (dark gray).

The average minimum distance between residue side chains and DMPC lipids (Figure S 7) shows that hydrophobic residues (e.g., V, F) generally exhibit shorter distances to the lipid bilayer compared to polar and charged residues. Phenylalanine (F) maintains consistently close contact with the membrane, indicating its dominant role in anchoring the peptide at the lipid interface. More subtle, sequence-dependent differences are observed for polar and charged residues. Lysine (K) residues in zi4F are often positioned further from the lipid bilayer, indicating weaker interaction with the membrane. In contrast, the corresponding glutamine (Q) residues in neg4F show more variable distances depending on their position in the sequence. This suggests that replacing K with Q reduces the tendency of these residues to remain away from the membrane. These differences are local and do not significantly affect the overall peptide–lipid interaction (Figure 4). Additionally, alanine (A) residues show position-dependent behavior, with distances varying depending on their neighboring residues, indicating a local sequence effect on peptide organization. Despite these local variations, the overall distance profiles remain similar between zi4F and neg4F, indicating that membrane association is primarily governed by conserved hydrophobic interactions.

Previous simulations have shown that changing the peptide sequence and charge pattern alters how helices arrange around the bilayer in nanodiscs and how they orient at the disc edge, highlighting the importance of specific residues and their electrostatic interactions.^26^ It was also shown that when the number of peptides is insufficient to fully cover the rim, the system relaxes by tilting the helices; this tilt allows the peptides to cover the edge more effectively and thereby stabilizes the nanodisc.

Consistent with these observations, the time evolution of the helix tilt angle in our simulations (Figure S 8) shows that both systems undergo an initial decrease in tilt during the early stages of the simulation, followed by stabilization. The tilt angle was defined as the angle between the vector connecting the Cα atoms of the N- and C-termini of the peptide and the membrane normal (z-axis). At the beginning of the simulation, tilt angles close to 90° indicate that the peptides are oriented approximately perpendicular to the membrane normal. Over time, the decrease in tilt reflects reorientation of the peptides toward a more aligned configuration relative to the membrane. The zi4F system maintains slightly higher tilt angles compared to neg4F throughout most of the trajectory. Despite this reorientation, visual inspection of the trajectories (Movies S 1–4) shows that parts of the nanodisc rim remain only partially covered by peptide, suggesting that additional peptide molecules would be required to fully shield the edge.

### Incorporation of Transmembrane Peptide

Polymer-, peptideor MSP nanodiscs are commonly used to study MPs. However, the expression, purification and functional characterization of MPs is complex and beyond the scope of this paper. As a proof-of-concept, we co-reconstituted a biotinylated transmembrane peptide into these PDC nanodiscs. The peptide sequence was adapted from the synaptobrevin transmembrane domain and included a helix breaker sequence (GSS) to add conformational flexibility (Figure 5a) and hydrophilic linker (EKPEK) to separate the transmembrane region from biotin and to enable efficient binding to streptavidin-modified quantum dots.

**Figure 5.**
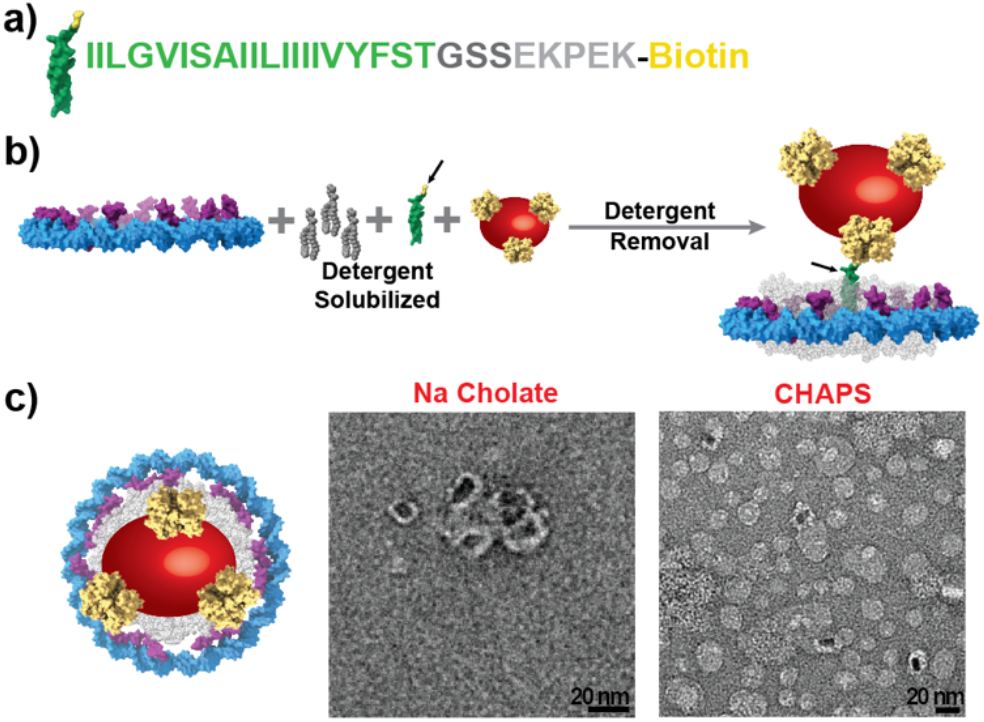
Incorporation of transmembrane peptide. a) Peptide sequence consisting of a hydrophobic transmembrane region, a flexible GSS linker, a hydrophilic linker (EKPEK), and biotin at the C-terminus. b) Co-reconstitution of detergent-solubilized lipids, the biotinylated trans-membrane peptide and streptavidin-modified quantum through detergent removal. B-c) Artistic representation of complex where with components drawn roughly to scale. c) TEM images of PDC nanodisc complexes with different detergents.

The modified quantum dots mimic bulky soluble domains or MPs and give a high contrast in TEM images. To synthesize the complexes, neg4F PDC minicircles were mixed with detergent-solubilized lipids and biotinylated transmembrane peptides, and streptavidin-modified quantum dots for several hours and then passed through a detergent removal column (Figure 5b) prior to TEM grid preparation. TEM images show QDs on top of the PDC lipid nanodiscs, demonstrating transmembrane peptide incorporation, thus indicating that the best efficacy was achieved for the sodium cholate system (Figure 5c). The outer diameter of these nanodisc complexes corresponds to 20±3 nm which is close to the calculated value of 18 nm for the outer diameter of an empty ds minicircle with 147 bp. Due to the large size of the quantum dots, the QD-modified PDC nano-discs did not migrate through the lane in PAGE (Figure S 9). A co-reconstitution experiment with CHAPS yielded lower transmembrane incorporation into the membranes compared to that of sodium cholate and more free lipids are seen in the background. Membrane protein reconstitution experiments typically require testing different detergents, too, to identify the most suitable system for the intended studies.

The solubility of the transmembrane peptide was tested in different detergents (sodium cholate, CHAPS and DTAB) and it was identified that it was soluble only in DTAB (Figure S 10a). We also tested the possibility of drying the methanol-solubilized transmembrane peptide along with the lipids during the lipid film preparation step but there was no incorporation of the peptide into the nanodiscs during the reconstitution step (Figure S 10b). The use of different detergents for solubilization of lipids and transmembrane peptides indicates that we can use these PDC minicircles for reconstitution of membrane proteins and lipids solubilized in different detergents.

## CONCLUSION AND OUTLOOK

In this work we introduced DNA-Peptide conjugate (PDC) nanodiscs as a new lipid bilayer mimetic that combines the advantages of structural DNA nanotechnology designs with the lipid bilayer stabilization abilities of popular amphiphilic nanodisc-forming peptides. Other than our previous DNA-based nanodiscs,^17,20^ PDC nanodiscs did not require any cationic detergents or lipids in the lipid reconstitution. Furthermore, we co-reconstituted a biotinylated transmembrane peptide that was coupled to a large streptavidin-modified quantum dot to mimic incorporation of large membrane proteins in future studies in structural Biology. Due to the flexibility of DNA-based designing, the circumference of the DNA scaffold as well as the number or the density of peptide modifications can easily be adjusted if needed and incorporated into larger DNA origami designs if needed, making them powerful tools for membrane protein characterization including Cryo-EM.

## ASSOCIATED CONTENT

Detailed material and methods used in this work; Sequences of DNA used; Synthesis of single stranded minicircle; SUPPLEMENTARY TABLES AND FIGURES: Micrographs of dsDNA minicircles using different modes of imaging; Optimization of PDC oligonucleotide excess; Optimization of gel loading concentration of PDC minicircles using 4F-DNA conjugates; Effect of detergent on PDC minicircles post-folding; HPLC-SEC characterization of mod4F PDC nano-discs; Average minimum distance between peptide side chains and DMPC lipids obtained from molecular dynamics simulations; Time evolution of the average tilt angle (θ) of peptide segments relative to the membrane normal; . 5% Native PAGE of transmembrane incorporated zi4F PDC nanodiscs; Solubility of transmembrane peptide in different detergents; Selection of capping atoms for parameterization; Chemical structures of DMPC; Crude HPLC chromatograms of the three peptides involved in this study; SUPPLEMENTARY MOVIES: All-atom molecular dynamics simulation of the DNA nanodisc (zi4F) using the CHARMM36 force field – side and top view; All-atom molecular dynamics simulation of the DNA nanodisc (neg4F) using the CHARMM36 force field-side and top view.

## AUTHOR INFORMATION

### Author Contributions

Conception of study: T.L.S., P.S.P.

Peptide Synthesis: J.K., D.P.N.G.

Peptide-DNA Conjugates: S.C., P.S.P.

PDC minicircles and nanodiscs: P.S.P.

TEM Imaging: P.S.P.

AFM Imaging: S.C.

Molecular Dynamics Simulations: F.F.

Supervision, Funding: T.L.S.

First draft of manuscript: P.S.P., S.C., F.F.

Revision of manuscript: all authors

### Competing Financial Interests

TLS is co-inventor of a patent (EP3755796B1) for a related technology.

## ACKNOWLEDGMENT

This research was funded by a MIRA grant from the National Institutes of Health/National Institute of General Medical Sciences (5R35GM142706) and the National Science Foundation through an EAGER grant (NSF EAGER 2017845) to T.L.S.

## ABBREVIATIONS

DNA: deoxyribonucleic acid
PDC: peptide-DNA conjugate
MP: membrane protein
EM: electron microscopy
MSP: membrane scaffolding protein
Salipro: saposin-lipoprotein
SMA: styrene maleic acid
NMR: nuclear magnetic resonance
MD: molecular dynamics
PEG: polyethylene glycol
ssDNA: single-stranded DNA
dsDNA: double-stranded DNA
zi4F: zwitterionic 4F peptide
D: aspartic acid
E: glutamic acid
K: lysine
F: phenylalanine
W: tryptophan
neg4F: negatively charged 4F peptide
Q: glutamine
RP-HPLC: reverse-phase high pressure liquid chromatography
PAGE: polyacrylamide gel electrophoresis
STEM: scanning transmission electron microscopy
CHAPS: 3-((3-cholamidopropyl) dimethylammonio)-1-propanesulfonate
TEM: transmission electron microscopy
SEC: size exclusion chromatography
V: valine
A: alanine
G: glycine
S: serine
P: proline
QD: quantum dots
DTAB: Dodecyltrimethylammonium bromide
TCEP: Tris(2-carboxyethyl) phosphine
PBS: phosphate-buffered saline
DMPC: 1,2-dimyristoyl-sn-glycero-3-phosphocholine
HEPES: 4-(2-hydroxyethyl)-1-piperazineethanesulfonic acid
Na_2_SO_4_: sodium sulfate
MgSO_4_: magnesium sulfate
SDS: sodium dodecyl sulfate
MgCl_2_: magnesium chloride
TBE: tris-boric acid-EDTA
EDTA: Ethylenediaminetetraacetic acid
AFM: atomic force microscopy
GROMACS: groningen machine for chemical simulations
CHARMM: chemistry at harvard macromolecular mechanics
CHARMM-GUI: chemistry at harvard macromolecular mechanics graphical user interface
CGenFF: chemistry at harvard macro-molecular mechanics general force field
CUFIX: champaign-urbana non-bonded fix
LINCS: linear constraint solver
CGeNArate: coarse-grained model for generating atomistic de-oxyribonucleic acid structures
web: world wide web
fs: femtosecond
PME: particle mesh ewald
V-rescale: velocity rescaling
C-rescale: stochastic cell rescaling
kJ: kilojoule
MDAnalysis: molecular dynamics analysis
MDVWhole: molecular dynamics voxel-based whole
nm: nanometer
mol: mole
rad: radian
TIP3P: transferable intermolecular potential with 3 points
Cα: carbon alpha
Gromologist: A groningen machine for chemical simulations-oriented utility library for structure and topology manipulation .

