## Supplementary material for "Peptide-DNA Conjugates for Formation of Lipid Nanodiscs": Suppl File

### MATERIALS AND METHODS

All oligonucleotides were purchased from Integrated DNA Technologies.

**Peptide Synthesis**

Peptides were synthesized using microwave-assisted solid-phase peptide synthesis on a CEM Liberty Blue 2.0 instrument employing Fmoc-chemistry. Rink Amide ProTide resin (CEM) was used as the solid support, with amino acid activation carried out using *N,N*'-Diisopropylcarbodiimide and Oxyma in DMF, and piperidine as the base. Resin cleavage and sidechain deprotection were performed using a mixture of trifluoroacetic acid (87.5 % v/v), phenol (5 % v/v), triisopropyl silane (2.5 % v/v), and water (5 %, v/v) at room temperature for 2.5 h. The crude peptides were precipitated in diethyl ether, collected by vacuum filtration then analyzed using HPLC.

**Peptide-DNA Conjugation**

10 µL of a 1 mM solution of HPLC-purified 3' and 5' thiolated oligonucleotides (/5ThioMC6-D/TTTTTCACACTTTTTCACACT/3ThioMC3-D/) were mixed with 2 µL of 500 mM Tris(2-carboxyethyl) phosphine (TCEP) (Sigma Aldrich) (10X molar excess) and incubated at room temperature for 2 h to reduce the dithiol bonds. TCEP removal was done by Sephadex (G-25) spin columns, according to the manufacturer's protocol, and the purified DNA was collected in 40 µL of 1X PBS (Phosphate-Buffered Saline (Thermo Scientific)) and was immediately added to 5 µL of 50 mg/mL maleimide-modified peptide (5X molar excess) dissolved in dimethylformamide (Fisher Chemical). The mixture was vortexed well and incubated at 25 □C, 1000 rpm for 12 h in Eppendorf Thermomixer C. Post-reaction, the sample was diluted with ultrapure water to 100 µL and purified by reverse phase (Viva C4 5 µm, Restek) HPLC (Agilent 1100 Series) using an acetonitrile (Supelco) gradient of 3-100% over 30 min, at a flow rate of 1 mL/min at room temperature. The respective peaks were collected, dried in Eppendorf Vacufuge Plus and the pellet was redissolved in ultrapure water.

**Concentration of PDC Conjugates**

The 260 nm absorbance of the dissolved pellet was measured using Thermo Scientific Nanodrop OneC spectrophotometer and the molar concentration was estimated for a path length of 1 cm and extinction coefficient of 192,398 M-1cm-1 (DNA e estimated using IDT OligoAnalyzer tool and peptide e estimated using ExPASy ProtParam tool).

**Formation of ssDNA Minicircles**

The protocol for formation of ssDNA was adapted from Iric et al.[27]

**Annealing**

4 μM (2 nmol) of 5’ phosphorylated linear ssDNA strands were incubated with 80 μM (20 nmol, 10X) of the splint strands in 1X T4 DNA ligase buffer (New England Biolabs). The strands were annealed in Hettich thermomixer by heating the strands to 80 °C for 5 min, followed by a 15-min cooling ramp from 80 to 50 °C and then a 10-min cooling ramp from 50 to 25 °C.

**Ligation**

Then, 10,800 U (27 μL of 400,000 U/mL) of T4 DNA Ligase (New England Biolabs) was added to the annealed mixture along with 1X T4 DNA ligase buffer. It was incubated at ambient temperature for 1 h, and then at 30 °C for 1 h. Heat inactivation was carried out by incubating the mixture at 65 °C for 20 min.

**Exonuclease Digestion**

12,000 U (60 μL of 20,000 U/mL) of exonuclease I (*E. coli.)* (New England Biolabs) and 48,000 U (48 μL of 100,000 U/mL) of exonuclease III (*E. coli.*) (New England Biolabs) were added to the ligated mixture along with 1X Exonuclease I Buffer (New England Biolabs) and 1X NEBuffer 1 (New England Biolabs). The mixture was incubated at 37 °C for 2 h and then heat inactivated by incubating at 80 °C for 20 min.

**Affinity Column Purification**

Purification of the covalently circularized ssDNA minicircles was performed using Zymo Spin V columns (Zymo Research) according to manufacturer protocol. Briefly, to 900 μL of the digested mixture, 1800 μL (2X volume excess) of the oligo binding buffer (Zymo Research) and 7200 μL (9X volume excess) of pure ethanol (Sigma Aldrich) was added and vortexed well. Centrifugation was performed using Eppendorf Centrifuge 5425. The Zymo Spin V column was placed inside a 2 mL VWR centrifuge tube and 800 μL of the sample mixture was added and centrifuged at 10,000 rcf for 30 s. The flow through was discarded and this step was repeated until all the sample mixture volume was exhausted. Then, 800 μL of the DNA wash buffer (Zymo Research) was added to the column and centrifuged at 10,000 rcf for 30 s. The flow through was discarded and the column was centrifuged at maximum rcf for 1 min to remove any residual buffer. The column was then transferred to a new 2 mL VWR centrifuge tube for sample elution. 800 μL of ultrapure water was added to the column and the sample was eluted by centrifuging at 10,000 rcf for 30 s.

**Concentration of ssDNA Minicircles**

The absorbance of the eluted ssDNA minicircles was measured at 260 nm on a Thermo Scientific Nanodrop One$^{C}$ spectrophotometer and the molar concentration was estimated for a path length of 1 cm and extinction coefficient of 1,608,800 $M^{-1}cm^{-1}$ (estimated using the IDT OligoAnalyzer tool).

**10% Urea PAGE**

10% Urea polyacrylamide gels were cast using Bio-Rad glass plates and gel casting setup. Gel runs were performed using Bio-Rad MiniPROTEAN® Tetracell system connected to the Power-Pac Basic power supply. The hand-cast gels are made up of 10% acrylamide/bisacrylamide (19:1) (VWR), 8 M urea (VWR), 0.1% N,N,N′,N′-Tetramethylethylenediamine (TEMED) (Bio-Rad) and 0.1% ammonium persulfate (APS) (VWR). Typically, 0.5 pmol of the sample was mixed with 6 µL of 2X urea loading dye (50% formamide (Beantown Chemical) and 10 mM NaOH (VWR)) and the volume was made up to 12 µL using 1X TE (5 mM Tris (VWR) and 0.5 mM ethylenediaminetetraacetic acid (EDTA) (VWR) at pH 8) buffer. The gels were run in heated 1X TBE (90 mM Tris-borate (Boric Acid (VWR)) at pH 8.3) and 2 mM EDTA) buffer in a heated water bath (approximately 60 °C) for 40-45 min. The gels were post-stained in 20 mL of 1X Sybr™Gold DNA gel stain (Invitrogen) for 5 min and were imaged using a Cytiva Typhoon FLA9000 gel imager with a Sybr™Gold Emission (LPB ≥ 510 nm) filter.

**<u>Formation of DNA minicircles</u>**

**Folding of dsDNA Minicircles**

100 pmol of the ssDNA minicircles (final concentration = 2.1 µM) were mixed with 1,050 pmol (1.5X per site) of complimentary oligonucleotides in 1X folding buffer (1X TE buffer containing 5 mM $MgCl_2$ (VWR)). The minicircles were annealed in the Bio-Rad C1000 Touch™ thermocycler by incubating the strands at 50 °C for 2 min and then cooling it down to 25 °C at the rate of -1 °C/min.

**5% Native PAGE**

5% Native polyacrylamide gels were cast using using Bio-Rad glass plates and gel casting setup. Gel runs were performed using Bio-Rad MiniPROTEAN® Tetracell system connected to the PowerPac Basic power supply. The hand-cast gels are made up of 5% acrylamide/bisacrylamide (29:1) (MP Biomedicals), 5 mM $MgCl_2$, 0.1% TEMED and 0.1% APS in 1X TBE buffer. 2-5 pmol of the sample was mixed with 4 µL of 6X no SDS loading dye (2.5% Ficoll®-400, 10 mM

EDTA, 3.3 mM Tris-HCl, 0.02% Dye 1 and 0.001% Dye 2 at pH 8) (New England Biolabs) and the volume was made up to 24 μL using 1X folding buffer. The gels were run in pre-chilled 1X TBE buffer containing 5 mM $MgCl_2$ at 4 °C for 60-90 min. The gels were post-stained in 20 mL of 1X Sybr™Gold DNA gel stain for 5 min and were imaged using Cytiva Typhoon FLA9000 gel imager with a Sybr™Gold Emission (LPB ≥ 510 nm) filter.

**<u>Formation of DNA nanodiscs</u>**

**Lipid Film Preparation**

Each lipid film contains 250-500 nmol of lipids. 1,2-dimyristoyl-sn-glycero-3-phosphocholine (DMPC) powder (Avanti Lipids) was initially solubilized in 2:1 chloroform (OmniSolv, Supelco): methanol (VWR) mixture to make a stock solution of 25 mg/mL (=36.87 mM). 250-500 nmol of 25 mg/mL DMPC stock was carefully transferred to glass test tubes and the film was dried, over a heated water bath, using a continuous flow of nitrogen gas. The films were then placed in Bel-Art SP Scienceware vacuum dessicator overnight and were capped with nitrogen (Linde) prior to long-term storage at -20 °C.

**Formation of DNA Nanodiscs**

Just before use, the lipid films were solubilized in either 45 mM of sodium cholate (Sigma Aldrich) or 16 mM of CHAPS (Thermo Scientific) and 1X buffer H (50 mM HEPES (VWR), pH 7.3, 100 mM $Na_2SO_4$ (Sigma Aldrich) and 8 mM $MgSO_4$ (Sigma Aldrich)). 7.5-200 pmol of dsDNA minicircles (final concentration = 0.3 μM) were incubated with 3.375-90 nmol of detergent solubilized lipids (final concentration = 1.69 μM), 45 mM sodium cholate or 16 mM CHAPS in 1x buffer H overnight. The sample mixtures were then subject to detergent removal using 125 μL (for sample volumes below 30 μL) or 500 μL Pierce detergent removal columns according to the manufacturer's protocols. Centrifugation was performed using Eppendorf Centrifuge 5420. Briefly, the columns were placed inside a 2 mL VWR centrifuge tube and were centrifuged at 1000 rcf (for 125 μL) or 1500 rcf (for 500 μL) for 1 min to remove the storage buffer. The location of the bed was marked, and it was ensured that the location of the bed does not change in the forthcoming centrifugation steps. 100 μL (for 125 μL) or 300 μL (for 500 μL) of 1X Buffer H was added to the column and was centrifuged at 1000 rcf (for 125 μL) or 1500 rcf (for 500 μL) for 1 min and the step was repeated twice to equilibrate the column. The flow-through was discarded. The column was centrifuged at 1000 rcf (for 125 μL) or 1500 rcf (for 500 μL) for 2 min to remove residual buffer prior to sample loading. Finally, 25 μL (for 125 μL) or

100 μL (for 500 μL) of sample was added to the column and was incubated for 2 min at ambient temperature (25 °C). The column was transferred to a fresh 2 mL VWR centrifuge tube, and the samples were eluted out by centrifuging at 1000 rcf (for 125 μL) or 1500 rcf (for 500 μL) for 2 min.

**HPLC-SEC Characterization**

HPLC-SEC was performed using Superdex™ 200 Increase 10/300 GL column connected to Agilent Series 1100 HPLC system. 100 μL of 175 pmol of sample was injected and was eluted at ambient temperature, in 50 mM Tris, pH 8.8 containing 100 mM NaCl (VWR) at a flow rate of 0.5 mL/min.

**5% Native SDS PAGE**

5% Native-SDS polyacrylamide gels were cast using Bio-Rad glass plates and gel casting setup. Gel runs were performed using Bio-Rad MiniPROTEAN® Tetracell system connected to the PowerPac Basic power supply. The hand-cast gels are made up of 5% acrylamide/bisacrylamide (29:1), 5 mM $MgCl_2$, 0.05% sodium dodecyl sulfate (SDS) (TCI Chemicals), 0.1% TEMED and 0.1% APS in 1X TBE buffer. 5 pmol of the sample was mixed with 1.5 μL of 10X native SDS loading buffer (100 mM Tris-HCl, pH 8.0 containing 0.5% SDS and 50 mM $MgCl_2$) and the volume was made up to 15 μL using 1X folding buffer. The gels were run in pre-chilled 1X TBE buffer containing 5 mM $MgCl_2$ and 0.05% SDS at 4 °C for 90 min. Post-run, the gels were briefly washed in 1X TBE buffer and then were stained in 20 mL of 1X Sybr™Gold DNA gel stain in 1X TBE buffer for 5 min and were imaged using a Typhoon FLA9000 gel imager with a Sybr™Gold Emission (LPB ≥ 510 nm) filter.

**<u>Incorporation of Transmembrane Peptide bound to Quantum Dots into the PDC Nanodiscs</u>**

250 nmol lipid films were used for transmembrane peptide studies. Just before use, lipid films were solubilized in either 45 mM sodium cholate or 16 mM CHAPS in 1X buffer H. 7.5 pmol of dsDNA minicircles (final concentration = 0.3 μM) were incubated with 7.5 pmol of transmembrane peptide solubilized in 100 mM DTAB (Sigma Aldrich) (final concentration = 0.3 μM), 0.25 μL of QdotTM 655 streptavidin conjugate (Invitrogen), 3.375 nmol of detergent solubilized lipids (final concentration = 1.69 μM), in final concentrations of 45 mM sodium cholate or 16 mM CHAPS in 1x buffer H overnight. The sample mixtures were then subject to detergent removal using 125 μL Pierce detergent removal columns according to the manufacturer's protocol (as described under "formation of DNA nanodiscs").

### TEM Grid Preparation and EM Imaging

#### TEM Grid Preparation

Carbon-only 400 mesh copper grids (Electron Microscopy Sciences) were plasma treated for 30 s using PELCO easiGlow$^{TM}$ just before use. 10 µL of sample was applied to the grid for 5 min and was wicked away using a VWR light-duty tissue wipe. The grid was then transferred to a 10 µL drop of freshly activated 2% uranyl formate (Electron Microscopy Sciences) stain (activated using 1.25 µL of freshly prepared 1 M NaOH) solution for 30 s. Excess stain was wicked away using a VWR light-duty tissue wipe and the grid was quickly dried using a continuous flow of nitrogen.

#### TEM Imaging

Grids were imaged on 200 kV Tecnai G2 F20 microscope equipped with Gatan CCD camera and the images were analyzed using Fiji ImageJ image analysis software.

### AFM Sample Preparation and Imaging

#### AFM Sample Preparation

20 µL of 0.01% polyornithine (Sigma Aldrich) solution were added onto freshly cleaved mica (PELCO, Ted Pella) and incubated for 2 min. The mica surface was then washed with ~10 mL of ultrapure water and dried with a stream of nitrogen gas. 10 µL of 20 nM dsDNA minicircles were deposited onto the mica disc and incubated for 3–5 min. The solution was wicked away and the surface was washed twice with 1 mL of ultrapure water after which 60 µL of ultrapure water was added on to the sample surface. 20 µL of ultrapure water were also added onto the ScanAsyst Fluid+ tip (Bruker).

#### AFM Imaging

Imaging was performed in liquid using the peak force tapping mode in a highspeed Bruker Dimension Fastscan bio atomic force. Images were processed with the Nanoscope analysis software.

### Molecular Dynamics Simulations

#### Model Construction and Preparation

The circular DNA model used in this study was generated using the CGeNArate web server.[28] The peptide structures were predicted using the AlphaFold web server.[29] The complementary 21-mer oligonucleotide was modified on 5' and 3' ends with the peptide (zi4F or neg4F) through a parameterized linker. To describe this linkage, two modified thymine residues were introduced:

TO5, modified at the O5′ atom, and TO3, modified at the O3′ atom. These residues were generated by extending the standard thymine (.rtp) entry with linker atoms whose parameters were derived from the CHARMM General Force Field (CGenFF).[30–32] To parameterize the nonstandard nucleotide, capping groups were added to the linker fragments to complete the chemical structure during charge calculation. After parameterization, these capping atoms were removed to obtain the final residues used in the full system (Figure S 11). Missing bonded or dihedral terms were derived by analogy using the Gromologist toolkit.[33] A pre-equilibrated DMPC bilayer was generated using CHARMM-GUI.[34–41] and combined with the DNA nanodisc. Lipids outside the DNA boundary or within 2.2 Å of the nanodisc were removed, resulting in 392 lipids (neg4F) and 381 lipids (zi4F) and producing clash-free structures for subsequent energy minimization and equilibration. Model construction and visualization were carried out using Discovery Studio[42] and PyMOL.[43] MDAnalysis[44,45] was used for structural refinement and trajectory analysis. All molecular dynamics simulations employed the CHARMM force field[46–51] with CUFIX corrections.[52–54]

**Simulation Parameters and Conditions**

Since the CHARMM force field is not fully optimized for DNA, long simulations can result in deviations from the native helical structure. Piecewise linear/harmonic restraint potentials (bond type 10) with a force constant of 10,000 kJ $mol^{-1}$ $nm^{-2}$ were applied to maintain base-pair hydrogen-bond distances. The restraints were applied to the N1 atoms of adenine and guanine and the N3 atoms of cytosine and thymine, ensuring stable base-pair geometry and maintaining the overall integrity of the DNA double helix. To reduce the simulation box size and prevent the DNA nanodisc from drifting, flat-bottom position restraints with a layer geometry were applied to the phosphorus atoms of eight selected DNA residues. These restraints defined a 6.0 nm unrestrained slab (±3.0 nm) along the normal nanodisc(z-axis). Within this region, the DNA scaffold was permitted free and unbiased motion, while a harmonic force constant (1000 kJ $mol^{-1}$ $nm^{-2}$) was applied only when the atoms moved beyond the ±3.0 nm threshold from their reference coordinates. The final simulation box dimensions were set to 21 nm × 21 nm × 12 nm and a minimum distance of 1.2 nm was maintained between the DNA nanodisc and the box boundaries. After defining the box, the system was solvated with the TIP3P water model, the default water model used by the CHARMM force field. The total charge of the neg4F system was –364 e, and that of the Zi4F system was –308 e. Both systems were neutralized by adding the appropriate number of $Na^+$ ions, and additional $Na^+$ and $Cl^-$ ions were added to reach a salt concentration of

150 mM. Energy minimization was performed in two stages using the steepest descent algorithm to eliminate unfavorable atomic contacts. In the first stage, no bond constraints (e.g., LINCS) were applied, and the DNA was kept unrestrained, while the peptide heavy atoms, dihedral angles and the phosphorus (P) atoms of the lipids were restrained until the energy gradient converged below 1,000 kJ $mol^{-1}$ $nm^{-1}$. In the second stage, constraints were applied to bonds involving hydrogen atoms using the LINCS algorithm, consistent with the CHARMM36 force field and restraints were applied to the DNA and peptide backbones, while their side chains were released. The lipid dihedral angles and phosphorus atoms remained restrained during this step. After minimization, the clash-free and stabilized structure obtained from the second stage was used as the reference for positional restraints in the subsequent six-step equilibration protocol adapted from CHARMM-GUI nanodisc setups[39,55] During equilibration, the restraints were gradually released to allow the system to relax. Restraints were applied to the backbone and side-chain atoms of both the DNA and peptide components and dihedral angles and the phosphorus (P) atoms of the lipids. For the peptides, the backbone atoms included N, Cα and C, while for DNA, it comprised P, O2P, O1P, O5′, O3′, C5′, C4′, C3′, O4′, C2′, and C1′. All remaining non-hydrogen atoms were treated as side-chain atoms.

The equilibration protocol consisted of three initial 1 ns steps with a 1 fs time step, followed by two 5 ns steps and one 50 ns step with a 2 fs time step (totaling 63 ns). Position restraints on the DNA backbone, side chains, and lipids were gradually reduced (DNA backbone: 4000 → 50 kJ $mol^{-1}$ $nm^{-2}$; side chains: 2000 → 0 kJ $mol^{-1}$ $nm^{-2}$; lipids: 1000 → 0 kJ $mol^{-1}$ $nm^{-2}$), along with dihedral restraints (1000 → 0 kJ $mol^{-1}$ $rad^{-2}$). For DMPC, position restraints were applied to the phosphorus atom (P) along the membrane normal (z-axis), and dihedral restraints were applied to C1–C3–C2–O21 (Figure S 12).

Temperature was maintained at 310 K using the V-rescale thermostat,[56] and pressure was controlled isotropically at 1 bar using the C-rescale barostat.[57] Long-range electrostatic interactions were treated with the Particle Mesh Ewald (PME) method[58], while van der Waals interactions were smoothly switched to zero between 1.0 and 1.2 nm using the Verlet cutoff scheme. All bonds involving hydrogen atoms were constrained with the LINCS algorithm.[59] Trajectories were saved every 50 ps. The resulting trajectories were corrected using the MDVWhole algorithm[60] to remove periodic boundary artifacts. All molecular dynamics simulations were performed using GROMACS 2024.4[61–63] on the Ohio Supercomputer Center.[64]

**Analysis**

Distances between peptide residues and the lipid bilayer were calculated using the GROMACS gmx pairdist tool. Peptide residues (excluding DNA and backbone atoms) were selected as the analysis group, while selected DMPC lipid tail carbon atoms were defined as the reference group. Distances were computed between the center of mass of each peptide residue and the specified lipid atoms, using a cutoff of 2 nm. Tilt angles of peptide segments were calculated relative to the membrane normal (z-axis) using the GROMACS gmx gangle tool. For each peptide segment, two Cα atoms corresponding to selected residue pairs were used to define a vector along the peptide backbone. The angle between this vector and the z-axis was then computed over the trajectory. All plots were generated using in-house Python scripts.

| Type | DNA Sequence (5' to 3') |
|---|---|
| ssDNA minicircle (Linear ultramer) | /5PHOS/ TGT GAA AAA AGT GTG AAA AAGTGT GAA AAA AGT GTG AAA AAGTGT GAA AAA AGT GTG AAA AAGTGT GAA AAA AGT GTG AAA AAGTGT GAA AAA AGT GTG AAA AAGTGT GAA AAA AGT GTG AAA AAGTGT GAA AAA AGT GTG AAA AAG |
| ssDNA minicircle Ligation splint | CTTTTTTCACACTTTTTCAC |
| Oligonucleotide conjugated to Peptide (PDC staple) | /5ThioMC6-D/TTTTTCACAC-TTTTTCACACT/3ThioMC3-D/ |
| Unmodified Staple | TTTTTCACACTTTTTCACACT |

**Table S 1. Sequences of DNA used.**

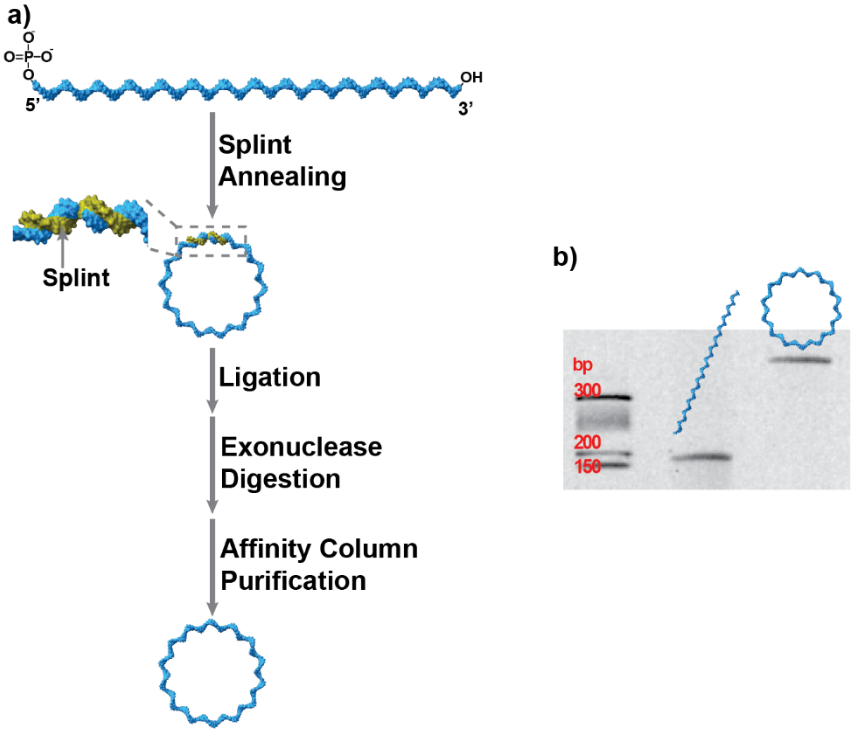


**Figure S 1. Synthesis of single stranded minicircle.** a) 5' phosphorylated strands were annealed to a complimentary 20mer oligonucleotide or "splint" and the ends were ligated using T4 DNA ligase. Residual linear fragments and splints were digested by exonucleases I and III and purified by affinity column purification (Zymo research). b) The formation of ssDNA minicircles was confirmed by denaturing PAGE. Note that ss minicircles migrate at a much slower rate than the linear strand.

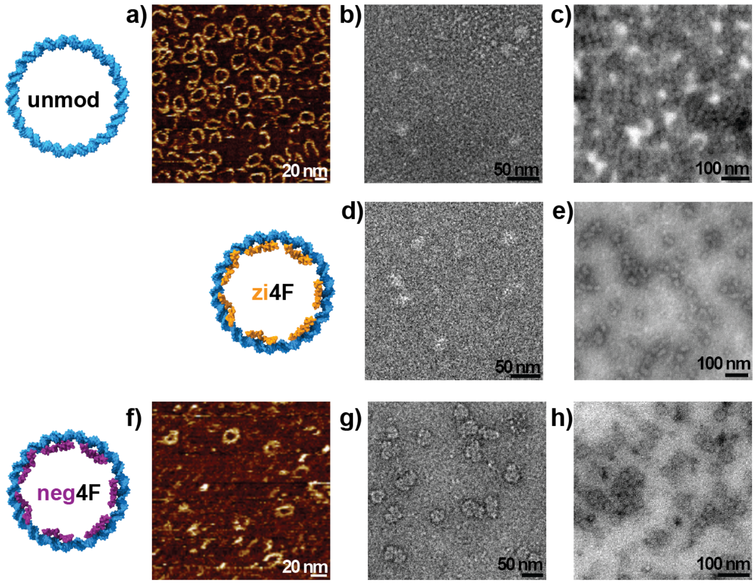


**Figure S 2. Micrographs of dsDNA minicircles using different modes of imaging.** a-c) Unmodified dsDNA minicircles imaged using AFM, TEM and STEM modes respectively, d-e) zi4F PDC minicircles imaged using TEM and STEM modes respectively and f-h) neg4F PDC minicircles imaged using AFM, TEM and STEM modes respectively. The minircircles clearly appear ring-like under AFM while the TEM images appear to be filled. We hypothesize that this could be an effect of stain crystals accumulating within the minicircle and thus making it appear filled. STEM images indicate the interior to be hollow, as evidenced by the dark minicircle borders and paler backgrounds. The zi4F minicircles also appear collapsed and this could be due to unfavorable electrostatic interactions between the peptide and the phosphates of DNA. This effect was reversed in the case of neg4F minicircles. Sizes of minicircles were measured to be 17±2 nm.

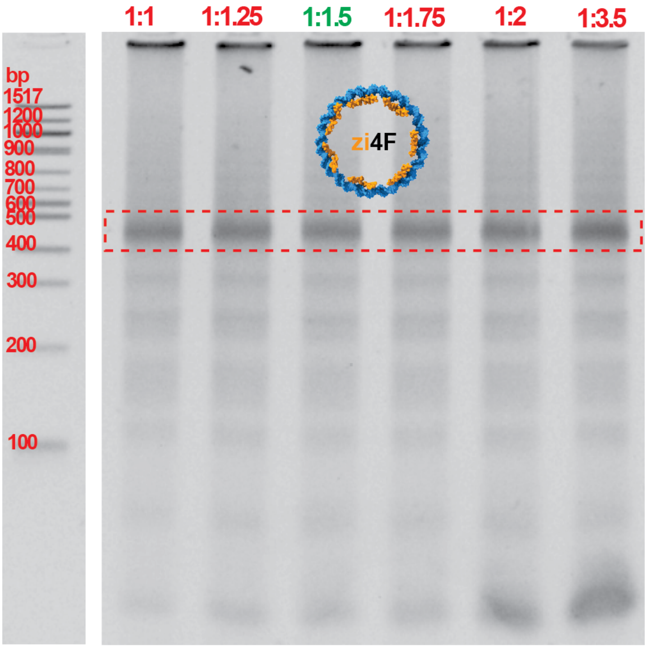


**Figure S 3. Optimization of PDC oligonucleotide excess.** Different ratios of ssDNA minicircle per binding site on short oligonucleotides were tested and 1:1.5 was a good compromise between high yields of dsDNA minicircles and avoiding a large excess of free conjugates, so that ultrafiltration could be avoided. Note that seven ss oligonucleotides can hybridize onto the repetitive ss minicircle, so that the molar excess of zi4F-PDC oligonucleotides was 10.5-fold (7*1.5).

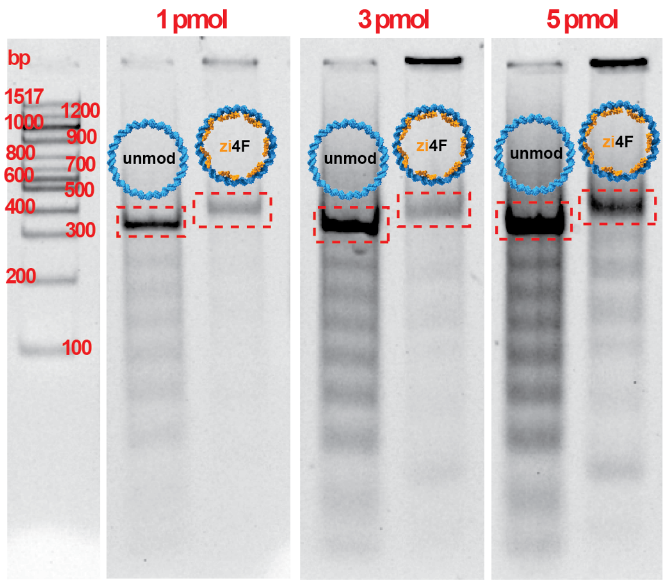


**Figure S 4. Optimization of gel loading concentration of PDC minicircles using 4F-DNA conjugates.** (a)Different ratios of ssDNA minicircle-to-short oligonucleotides were tested and 1:1.5 was optimal. This ratio meant that enough dsDNA minicircles were formed while keeping the excess free conjugates at a minimum, ensuring that the nanodiscs that will be formed would be those using the minicircles and not just the free peptide conjugates, while removing the need for ultrafiltration for purification of formed dsDNA minicircles. 5% native PAGE was used. (b) The intensity of the hanging lane for the PDC minicircles appeared to be dependent on the loading concentration, which indicates possible electrostatic aggregation occurring at the wells. This is hypothesized to be an effect of the presence of positive lysines in the peptide backbone binding to the negative phosphate DNA backbone. 5% Native PAGE was used.

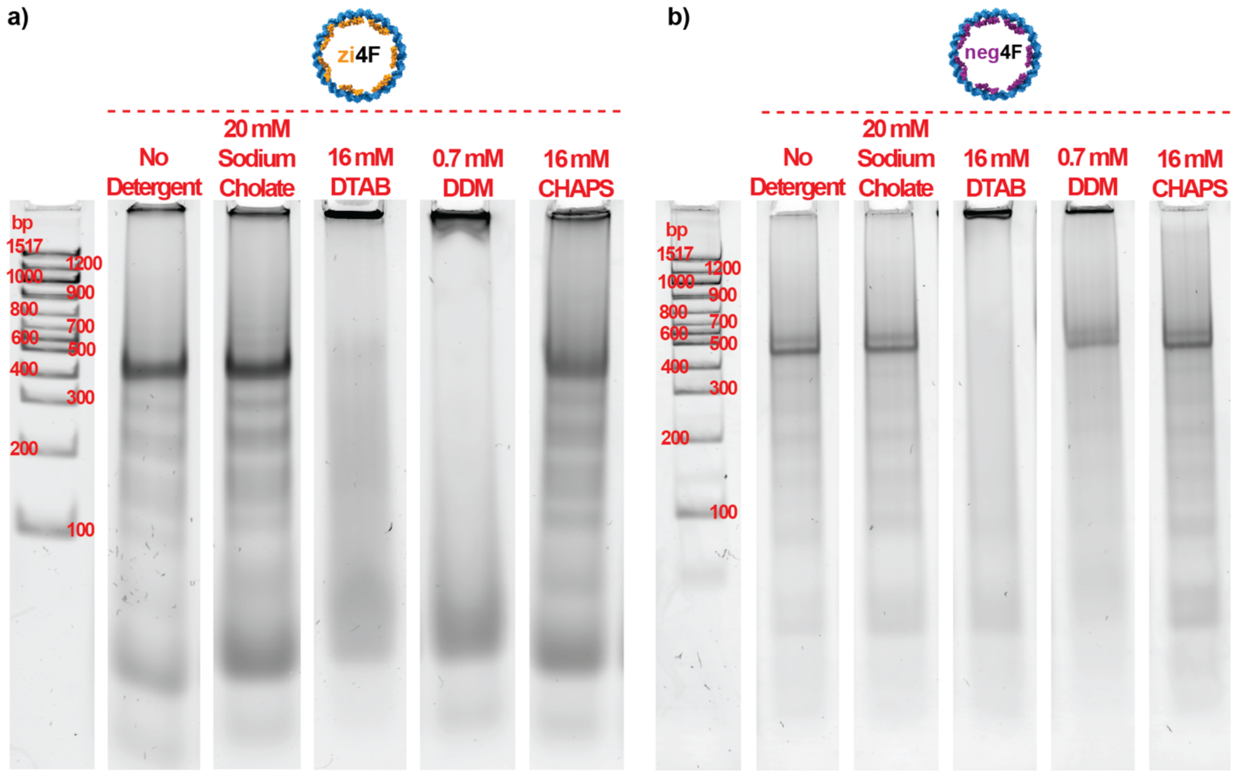


**Figure S 5. Effect of detergent on PDC minicircles post-folding**. As expected, DTAB, being a cationic detergent, electrostatically binds to the minicircle, rendering the overall charge of the system as neutral. This means that the minicircles no longer migrate into the gel. It is unclear whether there is structural collapse or not. In the case of DDM, it appears that there is some mode of interaction probably between the peptides and the detergent as the effect is much worse for the (a) zi4F PDC minicircles vs (b) the neg4F PDC minicircles. Based on this information, we decided to test the formation of nanodiscs using sodium cholate and CHAPS.

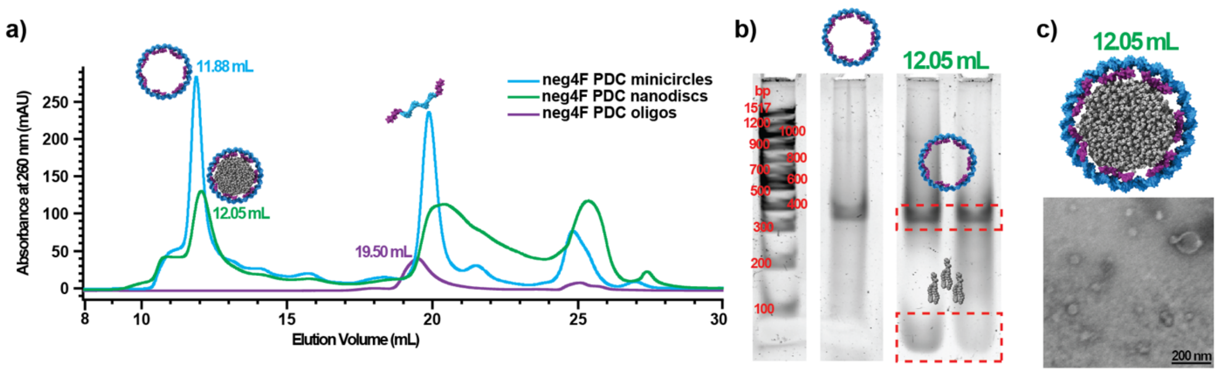


**Figure S 6. HPLC-SEC characterization of mod4F PDC nanodiscs.** a) HPLC-SEC profile of neg4F PDC oligos, minicircles and nanodiscs using DMPC:TfPC (99:1) lipids in sodium cholate. Detergent removal was performed using detergent removal spin columns prior to sample injection. Since the minicircles are covalently circularized, there is no elution peak shift between the minicircles and the nanodiscs. Still, the 12.05 mL peak, corresponding to that of the nanodiscs was collected and run on b) 5% Native-SDS PAGE. During the Native-SDS run, the SDS separates the PDC minicircles from the lipids and they appear as two separate bands (as indicated on the gel). When the same fraction was c) imaged using TEM, structures that looked like nanodiscs along with larger liposome-like structures were seen. The sizes of the nanodiscs were around 18 nm while the larger liposome-like strucutres were above 20 nm.

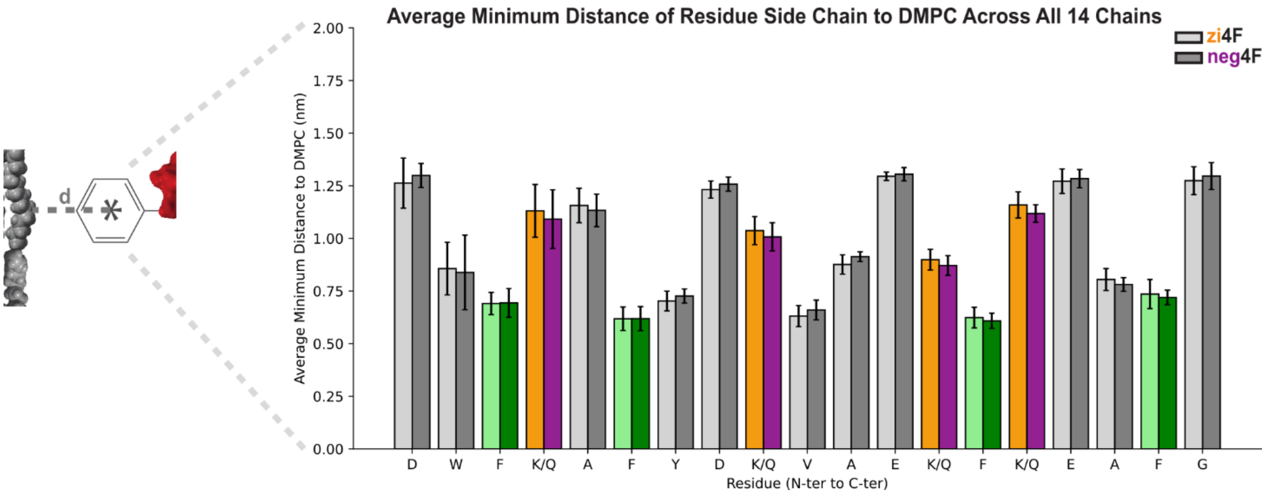


**Figure S 7. Average minimum distance between peptide side chains and DMPC lipids obtained from molecular dynamics simulations.** Values are shown for each residue and averaged over 14 peptide chains for the zi4F and neg4F systems.

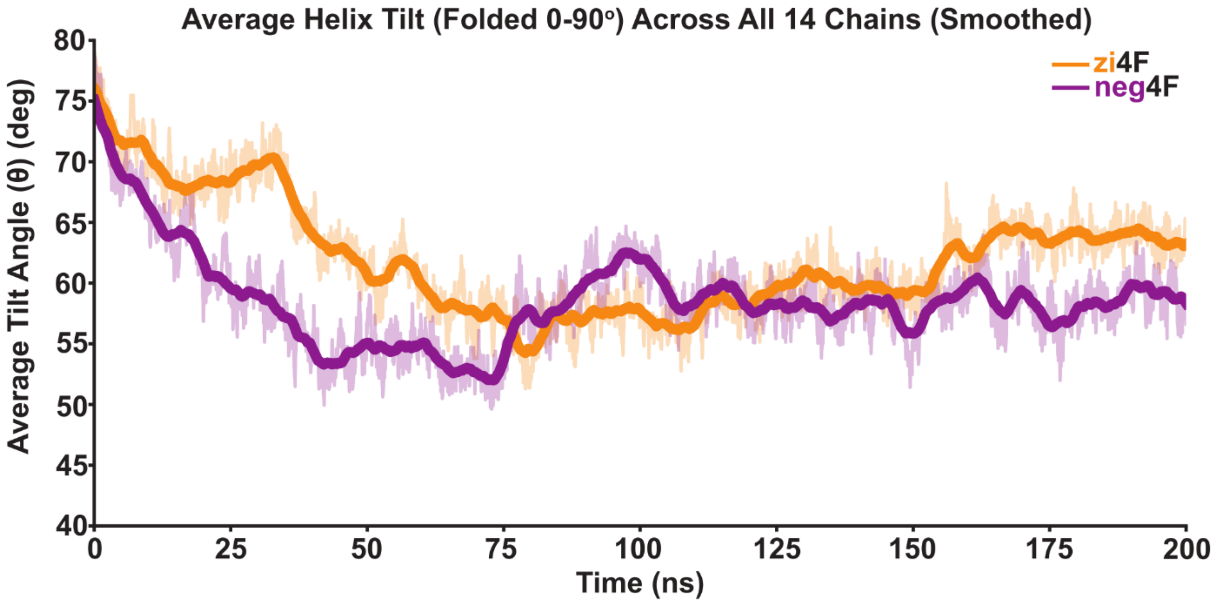


**Figure S 8. Time evolution of the average tilt angle (θ) of peptide segments relative to the membrane normal (z-axis)** over 200 ns for neg4F (purple) and zi4F (orange), showing differences in orientation and dynamic fluctuations between the systems.

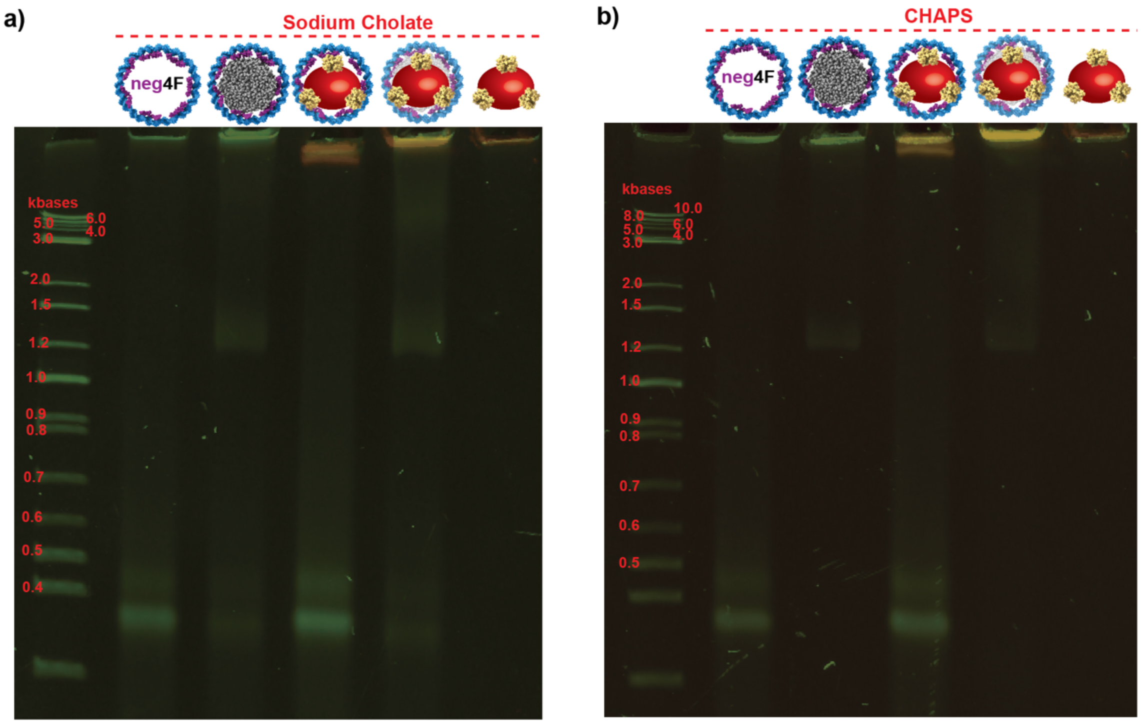


**Figure S 9. 5% Native PAGE of transmembrane incorporated zi4F PDC nanodiscs.** Green channel – Sybr$^{TM}$gold staining for DNA. Yellow channel – Quantum Dot (655 nm) signal. a) Sodium cholate solubilized lipids vs b) CHAPS solubilized lipids. Under both conditions, neg4F PDC minicircles and DTAB solubilized transmembrane peptides were used. The formed nanodiscs migrated at a higher rate when compared to the dsDNA minicircles. Due to the large size of the quantum dots, they are unable to migrate through the gel and stay at the well (as indicated by the hanging lane seen under QDs with only dsDNA minicircles, QDs with nanodiscs and only QDs). Thus, it is unclear from just PAGE characterization to ascertain whether the QD-peptide-nanodiscs complexes successfully formed or not.

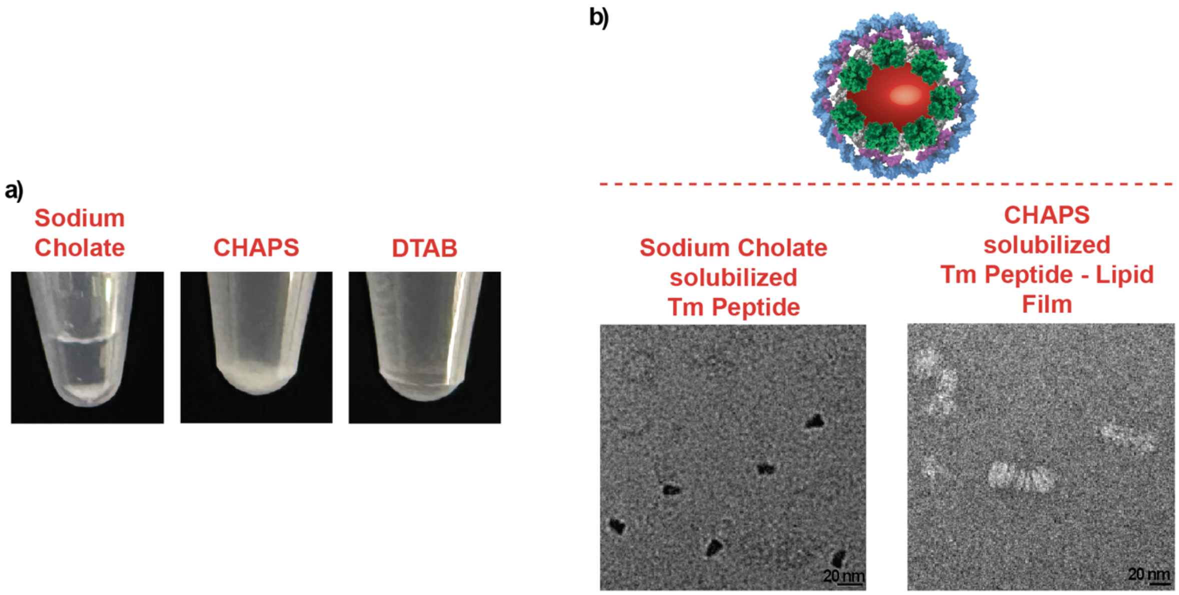


**Figure S 10. Solubility of transmembrane peptide in different detergents.** a) The transmembrane peptide was soluble only in DTAB and insoluble in other detergents such as sodium cholate and CHAPS. b) Adding the peptide to the lipid mixture directly during the lipid film preparation was unsuccessful.

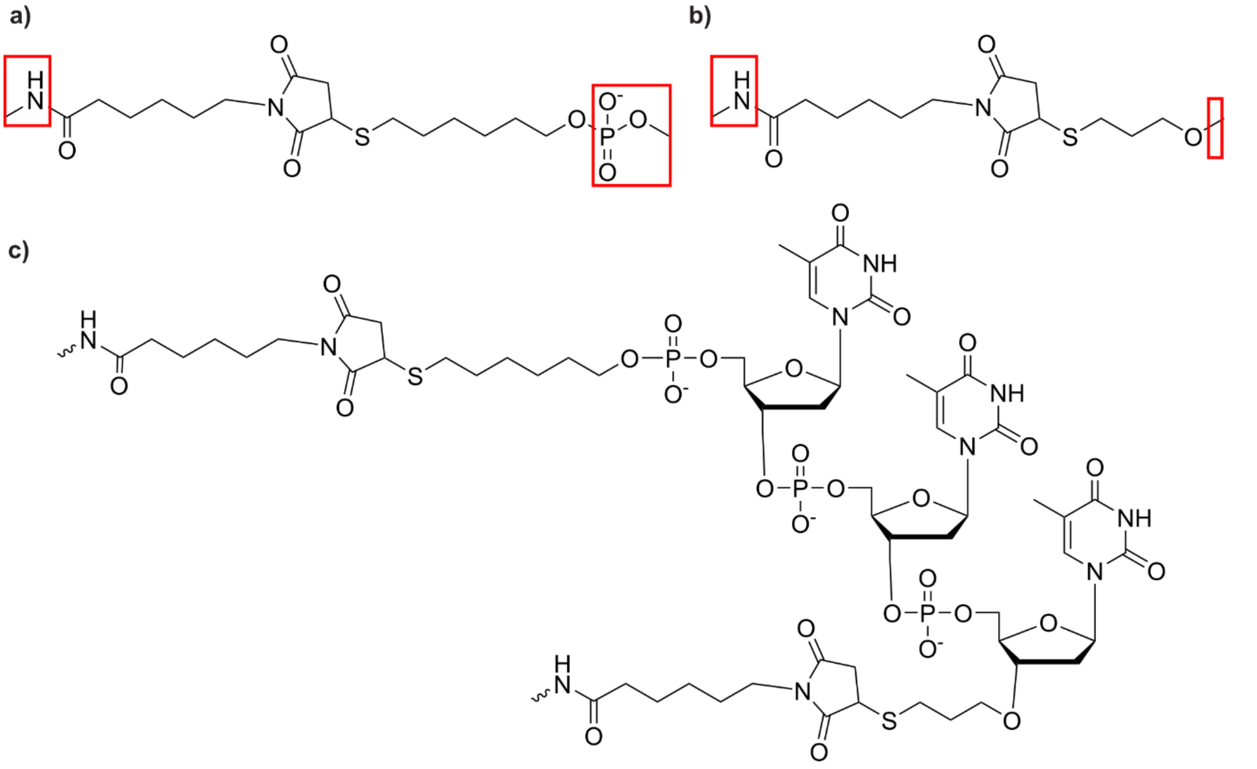


**Figure S 11. Selection of capping atoms for parameterization.** Terminal groups chosen as caps are highlighted with red boxes in the 5′ and 3′ thiol modifiers (a,b). (c) Final oligonucleotide structure after removal of the capping groups used for GROMACS parameterization. The first residue is labeled TO5, and the last residue is labeled TO3.

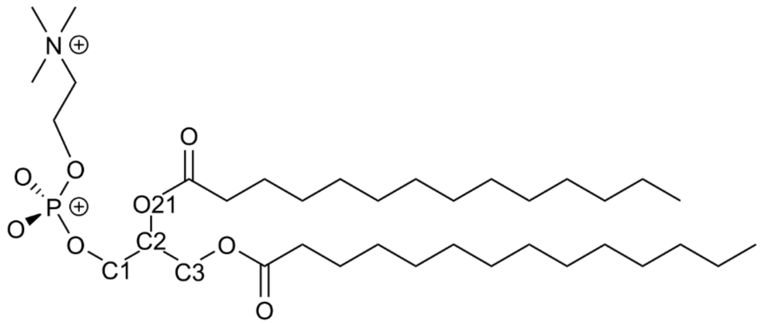


**Figure S 12. Chemical structures of DMPC.** The atoms used for applying dihedral restraints are (C1–C3–C2–O21). Position restraints were applied to the phosphorus atom (P) in DMPC.

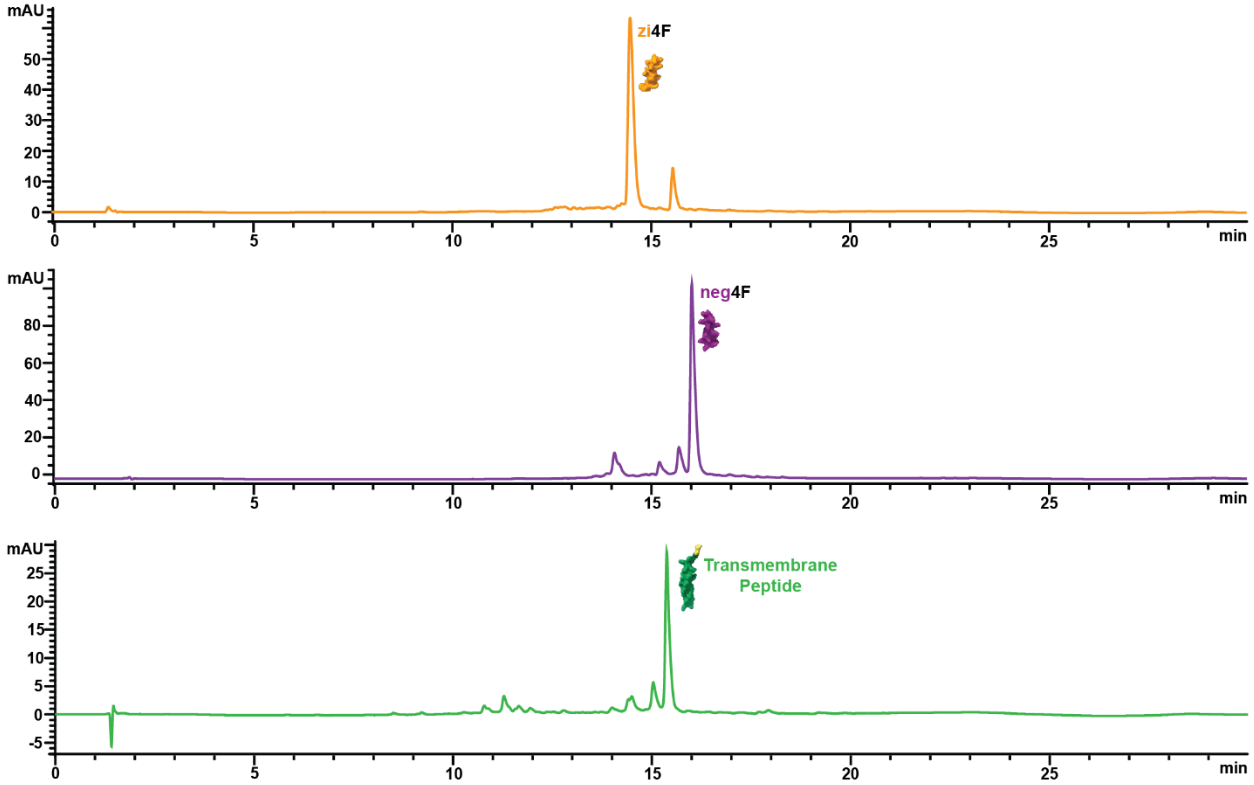


**Figure S 13. Crude HPLC chromatograms of the three peptides involved in this study.**

**Captions to Supplementary Movies**

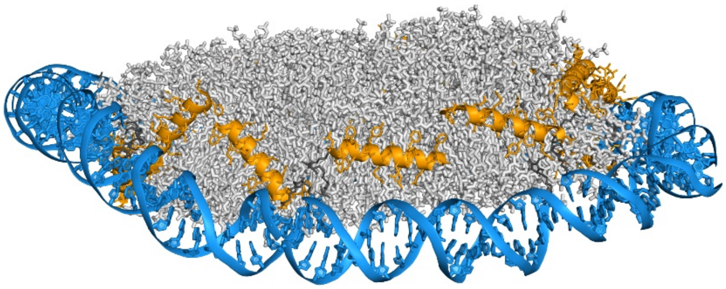

**Movie S 1. All-atom molecular dynamics simulation of the DNA nanodisc (zi4F) using the CHARMM36 force field. Side view** over 200 ns, showing interactions between the DNA scaffold (blue), peptides (orange), and lipid bilayer (gray). The snapshot shown is taken from the final frame at 200 ns.

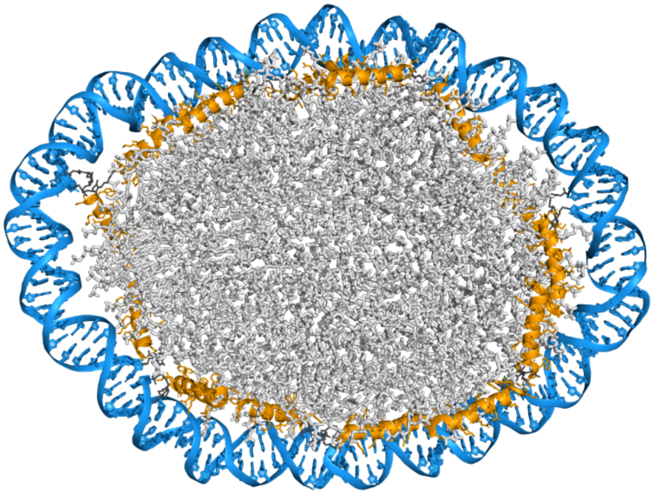

**Movie S 2. All-atom molecular dynamics simulation of the DNA nanodisc (zi4F) using the CHARMM36 force field. Top view** over 200 ns, showing interactions between the DNA scaffold (blue), peptides (orange), and lipid bilayer (gray). The snapshot shown is taken from the final frame at 200 ns.

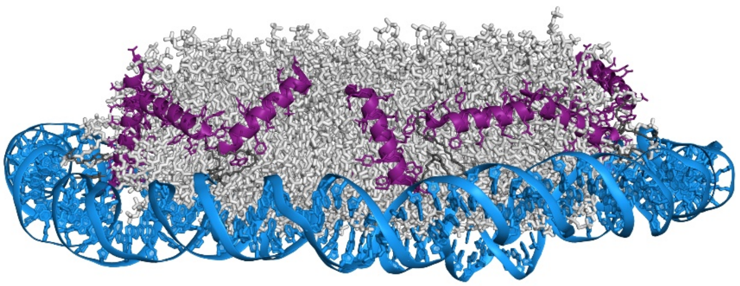

**Movie S 3. All-atom molecular dynamics simulation of the DNA nanodisc (neg4F) using the CHARMM36 force field. Side view** over 200 ns, showing interactions between the DNA scaffold (blue), peptides (magenta), and lipid bilayer (gray). The snapshot shown is taken from the final frame at 200 ns.

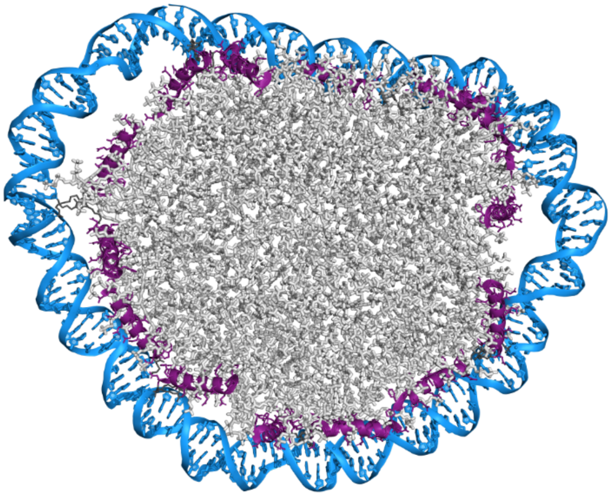

**Movie S 4. All-atom molecular dynamics simulation of the DNA nanodisc (neg4F) using the CHARMM36 force field. Top view** over 200 ns, showing interactions between the DNA scaffold (blue), peptides (magenta), and lipid bilayer (gray). The snapshot shown is taken from the final frame at 200 ns.
